# Common functional networks in the mouse brain revealed by multi-centre resting-state fMRI analysis

**DOI:** 10.1101/541060

**Authors:** Joanes Grandjean, Carola Canella, Cynthia Anckaerts, Gülebru Ayranci, Salma Bougacha, Thomas Bienert, David Buehlmann, Ludovico Coletta, Daniel Gallino, Natalia Gass, Clément M. Garin, Nachiket Abhay Nadkarni, Neele Hübner, Meltem Karatas, Yuji Komaki, Silke Kreitz, Francesca Mandino, Anna E. Mechling, Chika Sato, Katja Sauer, Disha Shah, Sandra Strobelt, Norio Takata, Isabel Wank, Tong Wu, Noriaki Yahata, Ling Yun Yeow, Yohan Yee, Ichio Aoki, M. Mallar Chakravarty, Wei-Tang Chang, Marc Dhenain, Dominik von Elverfeldt, Laura-Adela Harsan, Andreas Hess, Tianzi Jiang, Georgios A. Keliris, Jason P. Lerch, Hideyuki Okano, Markus Rudin, Alexander Sartorius, Annemie Van der Linden, Marleen Verhoye, Wolfgang Weber-Fahr, Nicole Wenderoth, Valerio Zerbi, Alessandro Gozzi

**Author notes:** Corresponding author Joanes Grandjean, PhD Singapore Bioimaging Consortium (SBIC) 11 Biopolis Way #01-02 Helios Building Singapore 138667.

## Abstract

Preclinical applications of resting-state functional magnetic resonance imaging (rsfMRI) offer the possibility to non-invasively probe whole-brain network dynamics and to investigate the determinants of altered network signatures observed in human studies. Mouse rsfMRI has been increasingly adopted by numerous laboratories world-wide. Here we describe a multi-centre comparison of 17 mouse rsfMRI datasets via a common image processing and analysis pipeline. Despite prominent cross-laboratory differences in equipment and imaging procedures, we report the reproducible identification of several large-scale resting-state networks (RSN), including a murine default-mode network, in the majority of datasets. A combination of factors was associated with enhanced reproducibility in functional connectivity parameter estimation, including animal handling procedures and equipment performance. Our work describes a set of representative RSNs in the mouse brain and highlights key experimental parameters that can critically guide the design and analysis of future rodent rsfMRI investigations.

## Introduction

The brain is the most complex organ, consisting of 86 billion neurons (Azevedo et al., 2009), each forming on average 7000 synapses. Approaching the complexity of the brain is rendered difficult due to the limited access to the tissue and the imperative for minimally invasive procedures in human subjects. Resting-state functional magnetic resonance imaging (rsfMRI) has gained attention within the human neuroimaging community due to the possibility to interrogate multiple resting-state networks (RSNs) in parallel with a relatively high spatial and temporal resolution (Biswal et al., 1995, 2010; Fox and Raichle, 2007). Functional connectivity (FC), i.e. the statistical dependence of two or more time series extracted from spatially defined regions in the brain (Friston, 2011), is the principal parameter estimated from rsfMRI studies. The importance of FC to neuroscience research can be understood through its widespread use to describe functional alterations in psychiatric and neurological disorders, e.g. for review (Buckner et al., 2008; Greicius, 2008). However, despite an extensive characterization of the functional endophenotype associated with diseased states, limitations with respect to invasiveness and terminal experiments generally preclude the establishment of detailed mechanisms in humans, as can be achieved with animal models.

Since its onset in 2011 (Jonckers et al., 2011), mouse rsfMRI has developed in a number of centres and has grown to become a routine method with a number of applications, reviewed in (Chuang and Nasrallah, 2017; Gozzi and Schwarz, 2016; Hoyer et al., 2014; Jonckers et al., 2015, 2013; Pan et al., 2015). Prominently, mouse rsfMRI has been used to investigate an extensive list of models, including Alzheimer’s disease (Grandjean et al., 2014b, 2016b, Shah et al., 2013, 2016c; Wiesmann et al., 2016; Zerbi et al., 2014), motor (DeSimone et al., 2016; Li et al., 2017), affective (Grandjean et al., 2016a), autism spectrum (Bertero et al., 2018; Haberl et al., 2015; Liska et al., 2018; Liska and Gozzi, 2016; Michetti et al., 2017; Sforazzini et al., 2016; Zerbi et al., 2018; Zhan et al., 2014), schizophrenia (Errico et al., 2015; Gass et al., 2016), pain (Buehlmann et al., 2018; Komaki et al., 2016), reward (Charbogne et al., 2017; Mechling et al., 2016), and demyelinating disorders (Hübner et al., 2017). Another application of mouse rsfMRI is the elucidation of large-scale functional alterations exerted by pharmacological agents (Razoux et al., 2013; Shah et al., 2016a, 2015). Finally, the method has been used to address fundamental questions. These include the investigation of the structural basis underlying FC (Bergmann et al., 2016; Grandjean et al., 2017b; Hübner et al., 2017; Schroeter et al., 2017; Sforazzini et al., 2016; Stafford et al., 2014), the nature of the dynamical event encoded in the resting-state signal (Belloy et al., 2018a, 2018b; Bukhari et al., 2018; Grandjean et al., 2017a; Sethi et al., 2017), as well as strain (Jonckers et al., 2011; Schroeter et al., 2017; Shah et al., 2016b), and the impact of sedation or awake conditions on the underlying signal and connectivity patterns (Bukhari et al., 2017; Grandjean et al., 2014a; Jonckers et al., 2014; Wu et al., 2017; Yoshida et al., 2016). This body of work obtained mainly over the past 5 years reflects the growth and interest into this modality as a translational tool to understand mechanisms underlying RSNs organisation in the healthy and diseased states, with the promise to highlight relevant targets in the drug development process and to advance fundamental knowledge in neuroscience.

Despite a growing interest in the field, rsfMRI studies in animals have been inherently difficult to compare. On top of centre-related contributions analogous to those observed in human studies (Jovicich et al., 2016), comparisons in rodents are further confounded by greater variability in preclinical equipment (e.g. field strength, hardware design), animal handling protocols and sedation regimens employed to control for motion and stress. Discrepancies between reports, such as the anatomical and spatial extent of a rodent homologue of the human default-mode network (DMN) (Becerra et al., 2011; Gozzi and Schwarz, 2016; Guilfoyle et al., 2013; Hübner et al., 2017; Liska et al., 2015; Lu et al., 2012; Sforazzini et al., 2014; Stafford et al., 2014; Upadhyay et al., 2011), or the organisation of murine RSNs (Jonckers et al., 2011), have stark consequences for the interpretations of the results. To meet a growing need to establish standards and points of comparison in rodent fMRI, we carried out a multi-centre comparison of mouse rsfMRI datasets. Multiple datasets representative of the local centre acquisitions were analysed with a common preprocessing pipeline and examined with seed-based analysis (SBA) and independent component analysis (ICA), two common brain mapping methods used to investigate RSNs. The aims of our work were to identify representative mouse RSNs, to establish a set of reference pre-processing and analytical steps and good-practices, and to highlight protocol requirements enabling more sensitive and specific FC detection in the mouse brain.

## Results

### Dataset description and preprocessing validation

A total of 17 datasets were included in this study. Dataset selection was restricted to 15 gradient-echo echo planar imaging acquired on C57Bl/6J mice, any gender, any age, any sedation protocol (**Supplementary table 1)**. Cortical signal-to-noise ratio (SNR) ranged from 17.04 to 448.56, while temporal SNR (tSNR) ranged from 8.11 to 112.68 (**Supplementary figure 1ab**). A comparison between SNR and tSNR indicated a positive association between the two measures (pearson’s r = 0.75, t = 18.30, df = 253, p = 2.2e-16). Due to the lack of orthogonality between the two factors, only SNR was considered in the remaining of the analysis. Mean framewise displacement (FWD) ranged 0.0025 mm to 0.15 mm (**Supplementary figure 1c**). A summary of representative estimated motion parameters is shown in the supplementary material (**Supplementary figure 2**). Each preprocessing output was visually inspected. Automatic brain extraction generated plausible brain masks. Normalisation was carried out to the Allen Institute for Brain Science (AIBS) template (**Supplementary figure 3**). Spatial coverage along the anterior-posterior axis varied across datasets. The following analysis is thus restricted to areas fully covered by all scans, corresponding to approximately 2.96 and -2.92 mm relative to Bregma. Moreover, distortions made it impossible to cover the amygdala region in full. No marked differences in the performance of each preprocessing steps were identified between datasets. The brain masked, spatially smoothed, temporally filtered, and normalized scans were further processed as follows.

### Vascular and ventricle signal regression enhances functional connectivity specificity

Denoising procedures are an integral step in all FC analyses relying on rsfMRI acquisitions. Nuisance signal originates from multiple sources, including physiological and equipment related noise (Murphy et al., 2013). No consensus exists both in human and rodent fMRI fields regarding optimal noise removal procedures. In this study the following six nuisance regression models were designed and compared with the aim to select one model based on objective criteria for the remaining of the analysis. The first nuisance model includes only motion correction and parameter regression (MC). Global signal regression (GSR) was added to the motion parameter in a second model. Signal from either white-matter (WM), ventricle (VEN), or vascular (VASC) masks (**Supplementary figure 4bcd**) were combined with motion parameters in additional regression models. Finally, based on results obtained with these approaches, a combination (VV) model including VEN and VASC signal regression was included for the comparison. The effectiveness of nuisance regression models and the specificity of the resulting networks at the subject level were assessed based on the outcome of a SBA using the anterior cingulate area (ACA, **Supplementary figure 4a**) as seed region. This seed was selected as a central node of the putative rodent DMN (Gozzi and Schwarz, 2016).

The statistical maps of the one-sample t-test across all individual maps following GSR (**Figure 1a**) indicated positive FC along rostro-caudal axis, through the ACA and extending to the retrosplenial area (RSP), with anti-correlations in adjacent primary somatosensory areas (SSp). Comparatively, in the VV nuisance model, a more extended network was revealed to include posterior parietal cortical areas (**Figure 1b**), while anti-correlations in the SSp did not reach statistical significance. To assess the specificity of the obtained functional networks, subject-level FC parameter (z-statistic) were extracted from ROIs located in the RSP and left SSp. The former was defined as a specific ROI, i.e. a ROI where positive FC is expected, while the latter was defined as a non-specific ROI, i.e. a ROI where low or negative FC is expected. The decision to consider these two areas as belonging to separable network systems reflects several lines of converging evidence: a) these regions are not linked by major white matter bundles or direct axonal projections in the mouse brain (Oh et al., 2014), b) they reflect separable electrophysiological signatures in mammals (Popa et al., 2009) c) they belong to separable functional communities (Liska et al., 2015) and are similarly characterized by the absence of significant positive correlation in corresponding human RSN (Fox et al., 2005).

**Figure 1.**
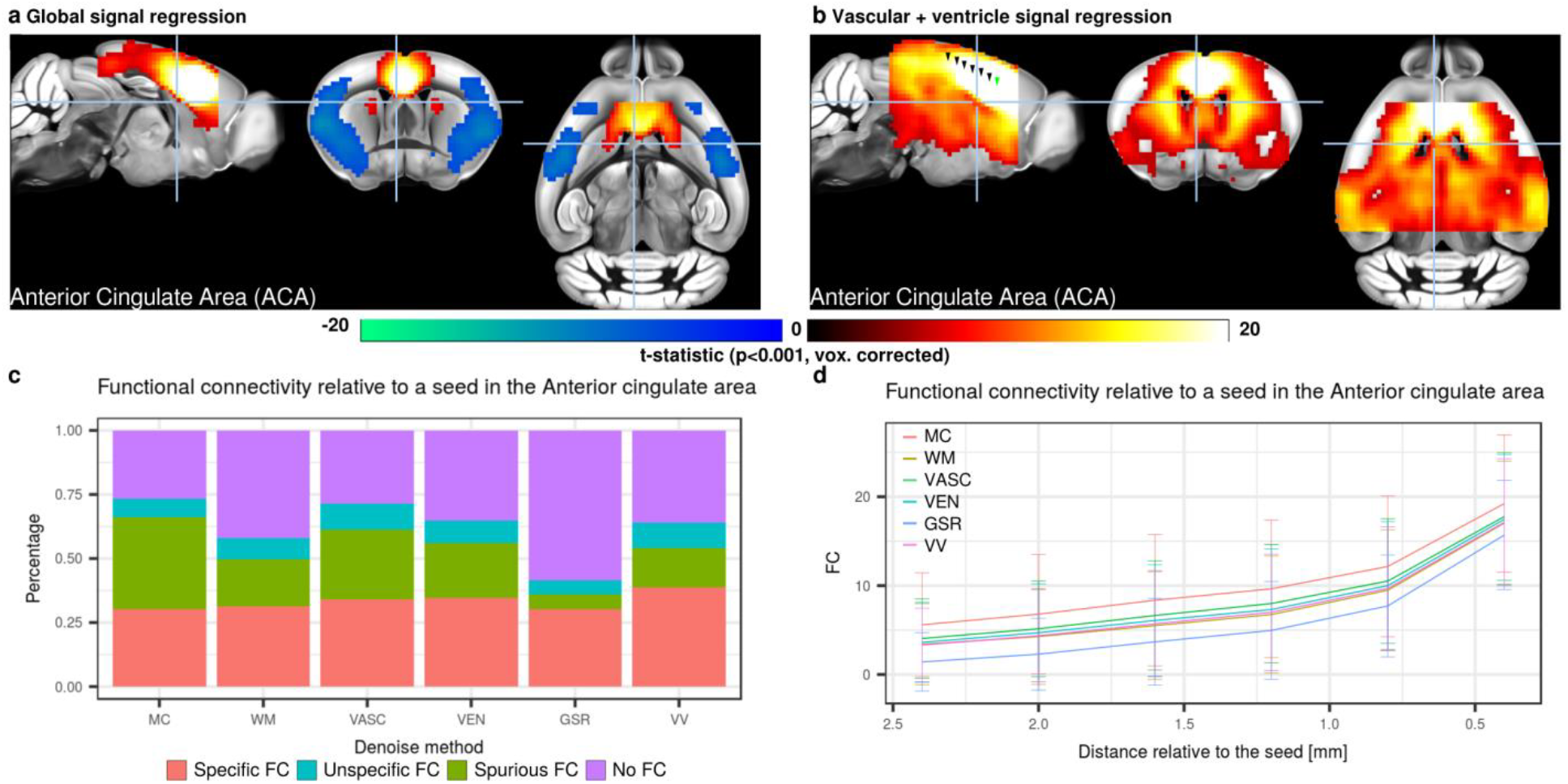
Denoising strategies and their impact on functional connectivity (FC) specificity. **a-b,** Seed-based analysis for a seed in the anterior cingulate area (ACA) following either global signal regression (GSR, **a**) or vascular+ventricle signal regression (VV, **b**). The spatial maps obtained lead to a set of regions for which the BOLD signals were positively associated to the BOLD signal of the ACA. These included the prefrontal cortex, retrosplenial area (RSP), dorsal striatum. Under VV, the connectivity profile extended to peri-hippocampal areas. Significant anti-correlation (negative t-statistic, blue) are also present in the primary somatosensory areas (SSp) under GSR but not VV condition. Individual scans were classified as presenting “Specific”, “Unspecific”, “Spurious”, or “No” FC relative to the ACA seed (**c**, see **Supplementary figure 5** for details). Comparison of each FC category depending on the denoising strategies revealed that motion correction and GSR lead to lowest percentage of “specific FC” at 30%, while that percentage was highest under VV condition (38%). FC as a function of distance to the ACA seed indicates comparable rate of decline between denoising strategies (**d**). Green arrowhead indicates the position of the ACA seed, black arrowheads indicate ROIs spaced 0.4 mm apart. Voxelwise corrected t-statistic for one-sample t-tests (p<0.001, corrected) are shown as a colour-coded overlay on the AIBS reference template. Descriptive statistics are shown as mean ± 1 standard deviation.

Detailed FC within the specific ROI for the GSR and VV nuisance model are shown as a function of FC within the corresponding non-specific ROI at the single-subject level (**Supplementary figure 5**). In the VV condition, 98/255 (i.e. 38%) of individual scans fell into the “specific FC” category while both MC and GSR reach lowest percentage (30%) of scans exhibiting “specific FC” relative to the ACA seed (**Figure 1c**). Out of the 98/255 scans categorised as presenting “specific FC” relative to the ACA seed, up to 14/15 scans originated from the same dataset (median = 6/15). Two datasets did not contain scans that met the definition. Correspondingly, the 98 scans were also unevenly distributed according to the different acquisition parameters, including field strength (4.7T N = 1/15, 7T N = 41/120, 9.4T N = 38/90, 11.7T = 18/30, X^2^ = 13.76, df = 3, p-value = 0.0032), coil type (room-temperature N = 26/105, cryoprobe N = 72/150, X^2^ = 13.13, df = 1, p-value = 0.00029), breathing condition (free-breathing N = 58/180, ventilated N = 40/75, X^2^ = 9.10, df = 1, p-value = 0.0026), and sedation condition (awake N = 7/15, isoflurane/halothane N = 18/90, medetomidine N = 26/75, medetomidine+isoflurane N = 47/75, X^2^ = 32.42, df = 3, p-value = 4.28e-07). Hence, scans presenting “specific FC” patterns were more often found in datasets acquired at higher field strengths, with cryoprobes, in ventilated animals, and under medetomidine+isoflurane combination sedation.

To test how FC is affected as a function of distance to the seed and nuisance model, FC in the ACA and RSP along the anterior -posterior axis was extracted (**Figure 1d**). Comparable rate of decrease was observed in all conditions, with GSR displaying an overall decrease of FC throughout. This is consistent with the overall decrease in FC induced by GSR relative to VV in the specificity analysis (**Supplementary figure 5**). In summary, the VV nuisance model enhanced specificity of SBA-derived DMN, as indicated by higher incidence of scans in the “specific FC” category. Based on this criterion, this nuisance model was used in all the subsequent analyses.

### Seed-based analysis identifies common and reproducible murine resting-state networks

We sought to identify common murine RSNs by means of SBA and to compare reproducibility across datasets. Seeds positioned in representative anatomical regions of the left hemisphere (**Supplementary figure 4a**) were used to reveal the spatial extent of previously described mouse resting-state networks. The seeds were selected to represent different cortical (somatosensory, motor, high order processing), as well as subcortical systems (striatum, hippocampal formation, thalamus). To obtain high-specificity and high-confidence group-level SBA maps, we first probed only the 98/255 scans listed as containing “specific FC” in the previous analysis. We next extended these analyses to include all the 255/255 scans (**Supplementary figure 7**). For datasets comparisons, all 15 scans from each dataset were included to reflect inter-dataset variability in the incidence maps.

All group-level SBA maps exhibited a strong bilateral and homotopic extension (**Figure 2a, Supplementary figure 6**). A seed in the ACA revealed a network involving the prefrontal cortex, RSP, dorsal striatum, dorsal thalamus and peri-hippocampal areas. This recapitulates anatomical features reminiscent of the human, primate and rat DMN (Gozzi and Schwarz, 2016; Hutchison and Everling, 2012; Sforazzini et al., 2014; Stafford et al., 2014). Comparable regions were observed with a seed in the RSP, a region evolutionarily related to the posterior cingulate cortex found in the human DMN (**Supplementary figure 6**). The anterior insular seed was found to co-activate with the dorsal cingulate and the amygdalar areas, corresponding to the putative rodent salience network (Gozzi and Schwarz, 2016), while the primary somatomotor region (MO) defined a previously described latero-cortical network that appears to be antagonistic to midline DMN regions, and that has been for this reason postulated to serve as a possible rodent homologous of the primate task-positive network (**Figure 2a**)(Liska et al., 2015; Sforazzini et al., 2014). Corresponding network across all scans (255/255) recapitulated features identified in the 98/255 scans listed as containing “specific FC”, but appeared to be characterized by much lower spatial specificity (**Supplementary figure 7**). Maps derived from individual datasets revealed that 70% (12/17) of the datasets presented the features listed above (**Figure 2b**, **Supplementary figure 8**). Incidence maps indicate, on a voxel basis, the percentage of the dataset presenting a significant FC. They confirmed the different extent of network detection in the different dataset. In summary, this analysis revealed the commonly shared spatial extent of mouse RSNs derived from SBA but also indicates that a small subset of the datasets failed to present these features with sufficient sensitivity or specificity.

**Figure 2.**
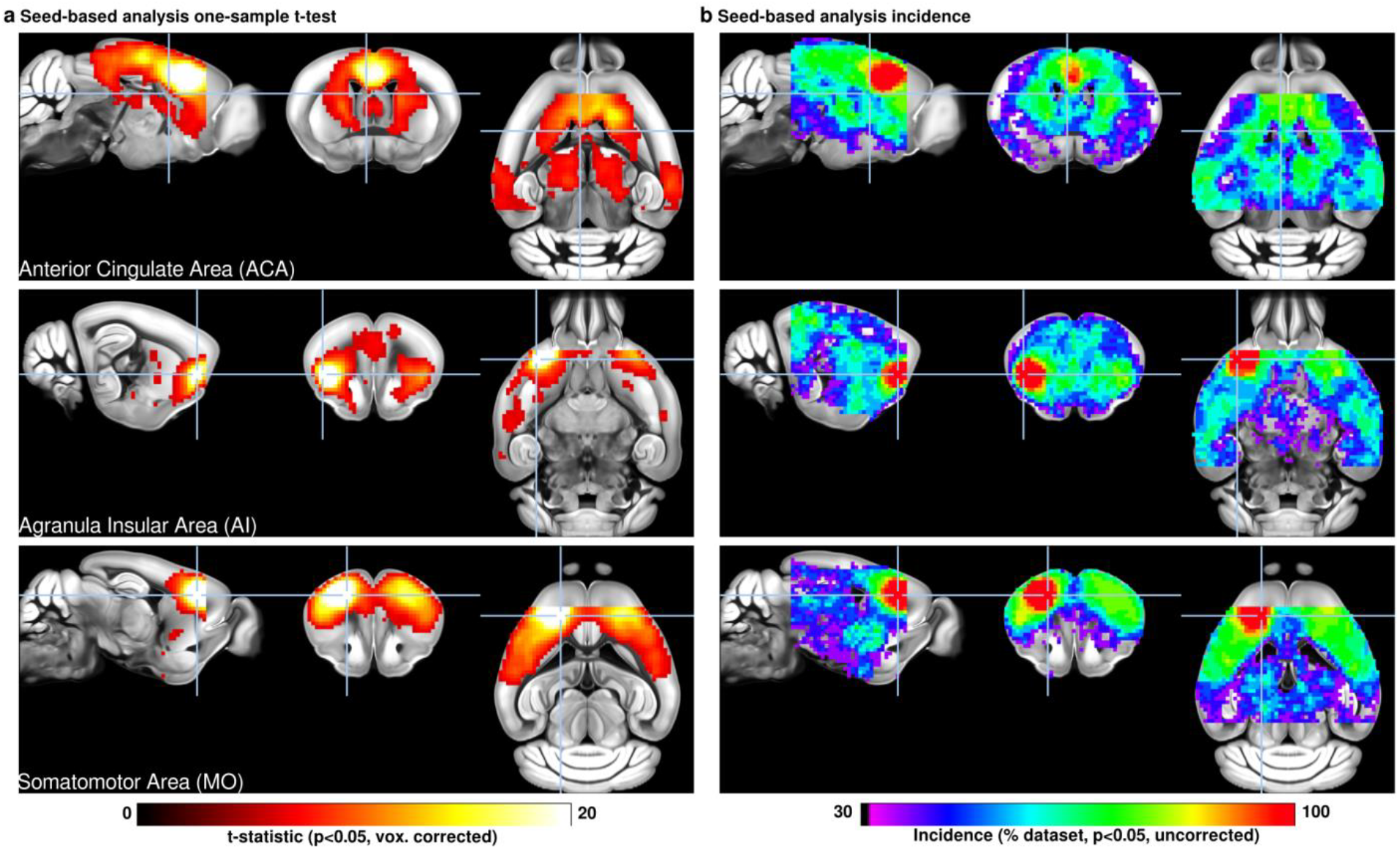
Seed-based analysis (SBA) for 3 selected seeds positioned on the left hemisphere. One-sample t-test maps of individual maps reveal the full extent of SBA-derived resting-state networks in the mouse brain across 98/255 scans that presented “specific FC” following vascular+ventricle signal regression. Functional connectivity (FC) relative to a seed located in the anterior cingulate area reveals the extent of the murine default-mode network, including the dorsal caudoputamen, dorsal thalamus, and peri-hippocampal areas. The seed in the insular area reveals significant FC in dorsal cingulate and amygdalar areas, corresponding to areas previously associated with the human salience network. Inter-hemispheric homotopic FC is found relative to the MO seed, together with lateral striatal FC. Incidence maps, indicating the percentage of dataset presenting significant FC in one-sample t-test (p<0.05, uncorrected), reveal that 12/17 of datasets recapitulated the features stated above. Out of these, 5 were not considered to overlap specifically (**Supplementary figure 6**). Voxelwise corrected t-statistic for one-sample t-tests and incidence maps are shown as a colour-coded overlay on the AIBS reference template.

### Sedation protocol and SNR affect connectivity strength

The datasets analysed here were acquired at varying field strengths, coil designs, EPI sequence parameters, animal handling, and with different anesthesia protocols, i.e. either awake or sedated states. Hence the acquisitions were not purposefully balanced to test for specific effects. To identify factors associated with FC strength, a simplified linear model was designed including the following explanatory factors: sedation and breathing conditions, SNR, and motion (mean framewise displacement). Limitations in orthogonality and representation of specific acquisition factors such as field strength, coil design, EPI sequence parameters, number of volumes, gender and age precluded a more extensive model.

Individual-level FC values (z-statistic) were extracted from SBA maps estimated from the ACA seed using a ROI located in the RSP and shown as a function of different acquisition parameters (**Figure 3**). Sedation protocol (F_(247, 251)_ = 18.29, p = 3.5e-13) and SNR (F_(247, 248)_ = 12.39, p = 5.1e-4) were significantly associated with FC, while the remaining factors, breathing condition (F_(247, 248)_ = 3.48, p = 0.063) and motion (F_(247, 248)_ = 0.082, p = 0.77) were not. The awake and medetomidine+isoflurane combination led to higher FC compared to the other two sedation categories. With respect to SNR, high FC values started to be observed at SNR > 50, suggesting that lower SNR may not be sufficient to detect relevant fluctuations. Interestingly, these effects were found consistently across the different ROI pairs considered (**Supplementary table 2**), thus confirming the importance of sedation conditions and SNR, and suggesting that breathing conditions impact mildly FC sensitivity.

**Figure 3.**
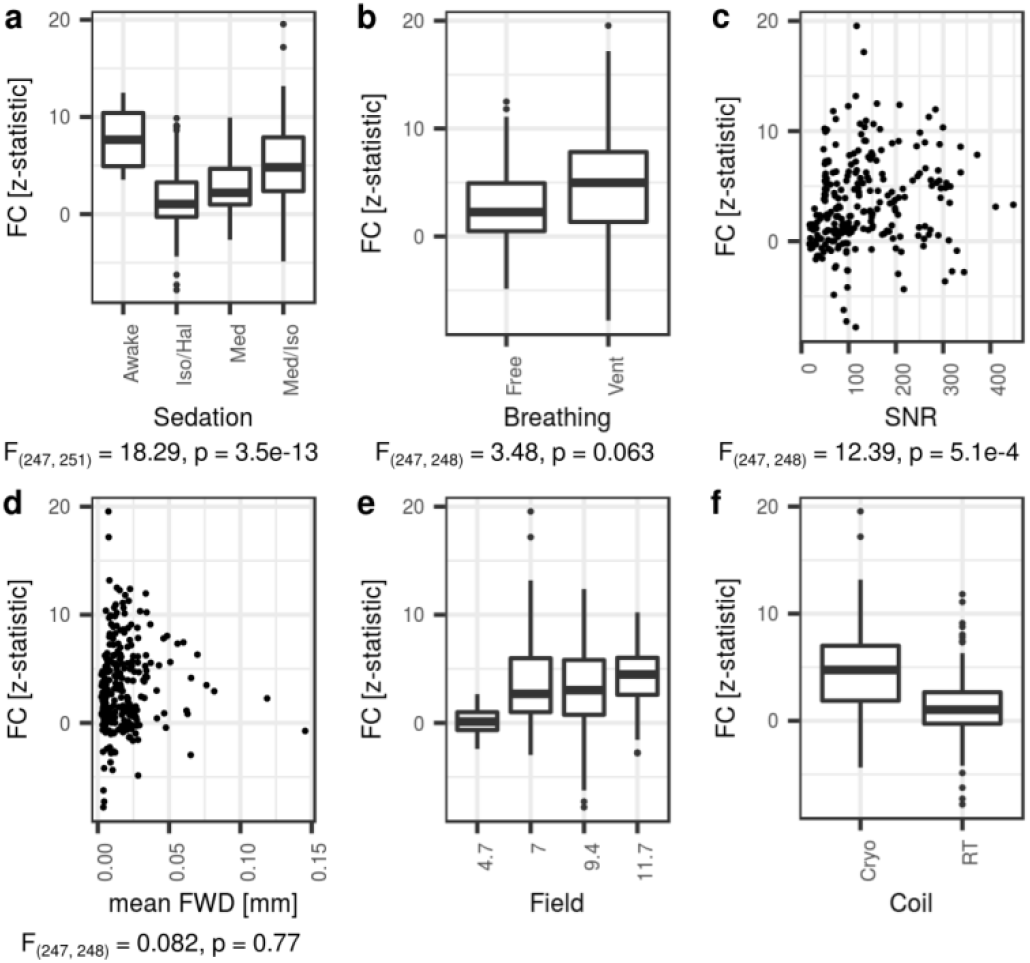
Functional connectivity (FC) in the retrosplenial cortex relative to a seed located in the anterior cingulate area, as a function of acquisition parameters. A statistically significant association was determined between sedation effect and FC (**a**, F_(247, =251)_ = 18.29, p = 3.5e-13) and between SNR and FC (**c**, F_(247,248)_ = 12.39, p = 5.1e-4). Neither breathing condition nor motion effects were significant with FC (**b**, **d**). Due to limitations in the representation of each level within a factor, coil design (**e**) and magnetic field (**f**) were omitted from the final statistical model. Free = Free-breathing, Vent = Mechanically ventilated, Cryo = cryoprobe, RT = room-temperature.

These animal handling conditions and sedation protocols highlighted here may not be applicable to all studies or laboratories due to local legislation, equipment availability, or technical knowledge. Distributions of FC values may hence provide useful reference points. Connectivity strength between the ACA and RSP, representing a central feature of the rodent DMN, reached z = 2.77, 5.71, and 10.46 at the 50^th^, 75^th^, and 95^th^ percentile respectively (Pearson’s r = 0.15, 0.26, 0.43, when SBA is carried out with a correlation analysis instead of a general linear model). Additional SBA parameter distributions are provided for other ROI pairs in **Supplementary table 2**. The parameters of the acquisitions featured in this analysis offer an objective criterion to evaluate and compare sensitivity to FC in a new dataset or in previous publications, insofar comparable metrics are available.

### Network-specific functional connectivity is found in all datasets

Evidence for robust distal FC could not be established in all datasets with SBA. To investigate the presence of network-specific FC also in datasets characterized by weaker long range connectivity, a dual regression combined with group-level ICA (drICA) approach was undertaken (Filippini et al., 2009). To obtain an enriched data-driven reference atlas, a group ICA atlas was generated out of the 98 “specific FC” scans selected in the SBA above, using 20 dimensions. The atlas revealed 9 cortical components (**Figure 4a, Supplementary figure 9, Supplementary table 3**), 5 overlapping with the latero-cortical network (somatomotor area (MO) and 4 SSp areas), 3 overlapping with elements of the DMN (prefrontal, cingulate/RSP, and temporal associative areas) and 1 overlapping with the insular area (AI). Additionally, 5 sub-cortical components were revealed, overlapping with the nucleus accumbens (ACB), caudoputamen (CP), pallidum (PAL), hippocampal region (HIP), and thalamus (TH) (**Supplementary figure 10**). The components recapitulate many of the features identified with SBA (**Figure 4b**), namely a strong emphasis on homotopic bilateral organization. The components identified here also presented strong similarities to a previous analysis (Zerbi et al., 2015). Due to uneven brain coverages across datasets, rostral and caudal RSNs could not be examined, including olfactory, auditory, and visual networks. To obtain individual-level representation of these components, a dual regression approach was implemented using the reference ICA identified above. These group-level ICA were used as masks to extract time series which were then regressed into individual scans using a general linear model. To investigate specificity relative to a DMN-related component, FC relative to the cingulate/RSP component was extracted from the ACA ROI (Specific ROI, z = 8.33, 14.41, and 22.32, 50^th^, 75^th^, and 95^th^ percentiles) and SSp ROI (Unspecific ROI). “Specific FC” was determined in 79% (201/255) of the scans, “Unspecific FC” in 16%, “Spurious FC” in 1.5%, and “No FC” in 3.1% (**Figure 4c**). “Specific FC” in 15/15 scans was determined in 2 datasets (Median = 12/15). The “Specific FC” category was also more evenly distributed relative to acquisition protocols and equipments: Field strength (4.7T N = 14/15, 7T N = 89/120, 9.4T N = 73/90, 11.7T N = 25/30, X^2^ = 4.01, df = 3, p-value = 0.25), coil type (room-temperature N = 88/105, cryoprobe N = 113/150, X^2^ = 2.17, df = 1, p-value = 0.14), breathing condition (free-breathing N = 138/180, ventilated N = 63/75, X^2^ = 1.29, df = 1, p-value = 0.25), and sedation condition (awake N = 13/15, isoflurane/halothane N = 65/90, medetomidine N = 55/75, medetomidine+isoflurane N = 68/75, X^2^ = 10.56, df = 3, p-value = 0.014). Importantly, statistical inference revealed that significant within-component FC could be established in 17/17 datasets for all 14 components (**Figure 4d**, **Supplementary figure 11, Supplementary figure 12**). This suggests that network-specific inferences can be probed in all rsfMRI datasets, and that drICA is a powerful approach enabling robust FC detection in all datasets, including those that may not robustly exhibit distal connectivity patterns.

**Figure 4.**
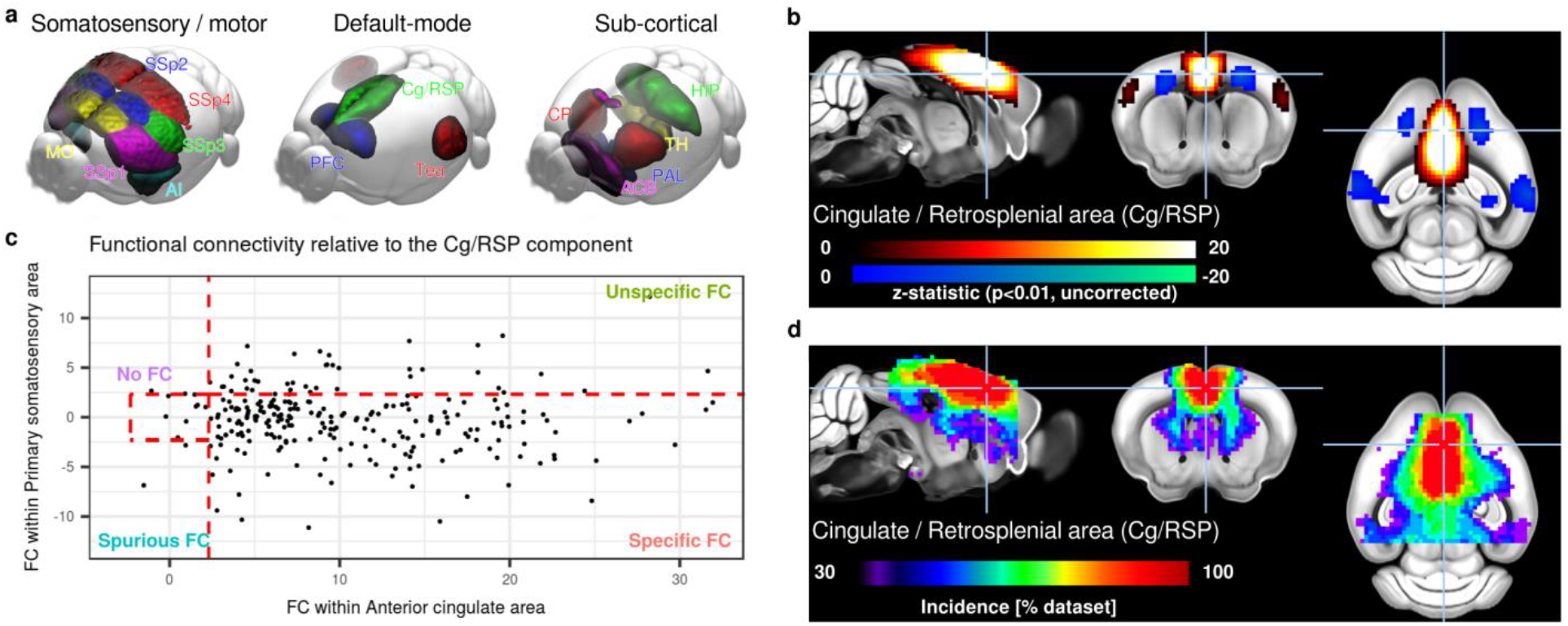
Group-level independent component analysis (ICA) estimated across 98/255 “specific FC” scans reveals canonical murine components (**a**). All components presented a marked bilateral organisation. Nine components were found to overlap principally with the isocortex including regions attributed to latero-cortical, salience and DMN networks by SBA, 3 components overlapped with the striatum, one with the hippocampal areas, and one with the thalamus. Detailed representations of the Cingulate / Retrosplenial area component (Cg/RSP **b**). Remaining components are presented in **Supplementary figure 9, 10**. FC relative to Cg/RSP is found specifically in the anterior cingulate area but not in the primary somatosensory in 79% of the individual scans following dual-regression (**c**). One-sample t-test within datasets indicates 100% of datasets presented significant FC (p < 0.05, uncorrected) within the Cg/RSP component. Incidence for the remaining components are presented in **Supplementary figure 11, 12**. AI = insular area, MO = somatomotor area, SSp = primary somatosensory area, PFC = prefrontal cortex, Cg/RSP = cingulate + retrosplenial area, Tea = temporal associative area, CP = caudoputamen, ACB = nucleus accumbens, PAL = pallidum, HIP = hippocampal region, TH = thalamus.

## Discussion

Rodent rsfMRI has been a growing research field in neuroscience over the past 10 years (Chuang and Nasrallah, 2017; Gozzi and Schwarz, 2016; Hoyer et al., 2014; Jonckers et al., 2015, 2013; Pan et al., 2015). The fast-paced development of the field has yielded a number of exciting results, yet the comparability of these findings remains unclear. The results presented here indicate that, despite major differences in cross-site equipment, scan conditions, sedation protocols and experience in the implementation of these procedures, mouse rsfMRI networks converge toward spatially defined motifs encompassing previously described neuroanatomical systems of the mouse brain. Importantly, we also highlight the possibility to use rsfMRI to probe distributed network systems of high translational relevance, including a rodent DMN, salience network, and latero-cortical network. While not reliably identified in all datasets and scan conditions, these large-scale networks were found to colocalize into well delineated boundaries in the majority of scans and datasets respectively, recapitulating previous descriptions in rodents (Gozzi and Schwarz, 2016; Lu et al., 2012; Sforazzini et al., 2014; Stafford et al., 2014), monkeys (Hutchison and Everling, 2012) and humans (Buckner et al., 2008).

Interestingly, most (70%) of the datasets converged toward spatially defined common RSN when long-range FC relative to a seed was assessed. When the analysis was restricted to local connectivity, all datasets converged. These results indicate that group-level, or second-level inferences, may be assessed irrespective of acquisition protocol or animal handling procedures in all datasets using robust analysis strategies. At the subject level, “specific FC” relative to the DMN was found in 38% of the scans, indicating that first-level inference on long-range FC is within reaches in some, but not all datasets. Sedation and equipment performance leading to increased SNR were the major factors associated with both FC sensitivity and specificity, together with breathing conditions. Awake animals presented higher FC overall, however datasets acquired with medetomidine+isoflurane combination together with mechanical ventilation were associated with greater specificity within elements of the DMN. Importantly, the results converged irrespective of sedation or awake protocols. This underlines that all datasets should be examined with the same expectations and criteria to further enhance results comparability. Hence, the set of standards provided here (e.g. spatial maps and FC parameter distributions), will allow the calibration of future multi-centre projects and assist in designing meta-analysis and replication studies, the gold standards in evidence-based research.

In addition to acquisition procedures, the adoption of analysis standards must be encouraged. A MRI template (Dorr et al., 2008) transformed into the AIBS standard space provides a common space that extends beyond animal MRI studies, including the seamless implementation of AIBS resources (Bergmann et al., 2016; Grandjean et al., 2017b; Oh et al., 2014; Richiardi et al., 2015; Stafford et al., 2014). Moreover, analysis based on robust methods (Zuo and Xing, 2014), such as drICA (Filippini et al., 2009), together with considerations for statistical analysis (Eklund et al., 2016), and sharing datasets on online repositories (Nichols et al., 2017) provide a comprehensive evidence-based roadmap to improve the comparability of acquisitions carried out between centres and enhance the robustness and reproducibility of future results. In particular, all the dataset analyzed in the context of this study will be shared and therefore provide references for scientists developing customized rsfMRI protocols.

Several major limitations within this study should be acknowledged. First and foremost, the lack of consensus quality assurance parameters for the estimation of FC led us to devise a strategy to examine FC specificity. Because this study grouped together a set of existing scans, factors were not entirely orthogonal and it was not possible to model a number of potentially relevant effects impacting FC metrics, such as specific sequence parameters (e.g. number of volumes), as well as biologically relevant factors including sex, age, and mouse strain. Finally, lack of distal FC in some datasets could not be attributed to specific animal handling protocols or equipment performance. This indicates that additional experimental factors not considered here may be better predictors estimating this particular kind of FC. For example, the implementation of procedures to control the arterial level of carbon dioxide may be critical to prevent hypercapnic conditions, a feature that is associated with reduced FC connectivity (Biswal et al., 1997) and that is often observed in freely breathing anesthetized rodents. Despite these limitations, the work presented here is likely to enhance the true scientific value of mouse rsfMRI by establishing standards and how to attain them. With these, the field is set to meet its goals toward the establishments and understanding of the cellular and molecular mechanisms of large-scale brain functional reorganisation in the healthy and diseased brain.

### Material and methods Comparison dataset acquisition

All animal experiments were carried out with explicit permits from local regulatory bodies. Seventeen datasets, consisting of 15 individual pre-acquired rsfMRI scans each, were acquired with parameters reflecting each centre standards. A summary of equipment, acquisition parameters, and animal handling procedures is listed in **Supplementary table 1**. Scans were acquired on dedicated Bruker magnets operating at 4.7T (N = 1 dataset), 7T (N = 8), 9.4T (N = 6), 11.7T (N = 2), with either room-temperature coils (N = 7) or cryoprobes (N = 10). Gradient-echo echo planar imaging (EPI) sequences were used to acquire all datasets, with repetition time (TR) ranging 1000 -2000 ms, echo time (TE) 10 -25 ms, and number of volume 150 -1000. Acquisitions were performed on awake (N = 1) or anesthetized C57Bl/6J mice (both male and female) with either isoflurane 1-1.25% (N = 5), halothane 0.75% (N = 1), medetomidine 0.1-0.4 mg/kg bolus and 0.2-0.8 mg/kg/h infusion (N = 5), or a combination of isoflurane 0.2-0.5% and medetomidine 0.05-0.3 mg/kg bolus and 0-0.1 mg/kg/h infusion (N = 5). Awake mice were fitted with a non-magnetic head implant to fix the heads to a compatible cradle (Yoshida et al., 2016). Animals were either freely-breathing (N = 12) or mechanically ventilated (N = 5). Datasets are publicly available in BIDS format on openneuro.org (project ID : Mouse_rest_multicentre, https://openneuro.org/datasets/ds001720).

### Data preprocessing

Volumes were analysed in their native resolution. Firstly, image axes were reoriented into LPI orientation (*3dresample*, Analysis of Functional NeuroImages, AFNI_16.1.26, https://afni.nimh.nih.gov) (Cox, 1996). Temporal spikes were removed (*3dDespike*), followed by motion correction (*3dvolreg*). Brain masks (*RATS_MM,* https://www.iibi.uiowa.edu*) (Oguz et al., 2014) were estimated on temporally averaged EPI volume (fslmaths*). Motion outliers were detected based on relative framewise displacement estimated during motion correction. Volumes with spikes or framewise displacement greater than 0.100 mm, corresponding to approximately 0.5 voxel of the average in-plane resolution, were labelled in a confound file to be excluded from later seed-based analysis and dual-regression. Linear affine parameters and nonlinear deformations with greedy SyN diffeomorphic transformation (*antsIntroduction.sh*) were estimated relative to a reference T2 MRI template (Dorr et al., 2008) registered into the AIBS Common Coordinate Framework (CCF v3, http://www.brain-map.org/) resampled to 0.200 mm^3^. Normalisation to AIBS space was carried out on brain masked EPI directly using ANTS (Advanced Normalization Tools, http://picsl.upenn.edu/software/ants/) (Avants et al., 2014, 2011). Anatomical scans corresponding to each EPI acquisition were not available in all cases. Despite this limitation, plausible registrations of murine EPI directly onto a T2 MRI template were rendered possible due to the relatively simple structure of the lissencephalic cerebrum and high EPI quality. Individual registered brain mask were multiplied (*fslmaths*) to obtain a study mask. The analysis was bounded within this study mask, i..e the brain areas covered by all individual scans. References to anatomical areas are made with respect to the AIBS atlas. All brain masks and registrations were visually inspected and considered plausible.

Six different denoising approaches were applied: i) 6 motion parameters regression (MC), or the following together with motion parameters, ii) white matter (WM), iii) ventricle (VEN), iv) vascular (VASC), v) vascular + ventricle (VV), or vi) global (GSR) signal regression. White matter and ventricle masks were adapted from the AIBS atlas (**Supplementary figure 4cd**), a vascular mask was obtained by averaging and thresholding hand-selected individual-level independent components registered to AIBS space (**Supplementary figure 4b**). Inverse transformations were applied to each mask. Average time series within masks were extracted (*fslmeants*) and regressed out (*fsl_regfilt*). Finally, spatial smoothing was applied with a isotropic 0.45 mm kernel (*3dBlurInMask*), and bandpass filtering was applied between 0.01 -0.1 Hz (*3dBandpass*). The smoothing kernel was selected to correspond approximately to 1.5 x voxel dimension of the lowest in-plane resolution. The bandpass filter was applied to all datasets to enhance comparability between datasets, despite indications that medetomidine leads to a shift in resting fluctuation frequencies (Grandjean et al., 2014a; Kalthoff et al., 2013; Paasonen et al., 2018). The denoised and filtered individual scans were normalised to AIBS reference space (*WarpTimeSeriesImageMultiTransform*).

Noise was estimated by extracting the signal standard deviation from manually defined regions-of-interest (ROIs) in the upper corners of at least 3 slices, carefully avoiding ghosting artefacts or tissues (brain or otherwise). Mean signal was extracted from the 20^th^ acquisition volume using a cortical mask spanning over the whole isocortex (defined by AIBS atlas) and registered in individual spaces to estimate signal-to-noise ratio (SNR). The same cortical mask was used to extract standard deviation of temporal signals to estimate temporal SNR (tSNR).

### Seed-based analysis and independent component analysis

Seeds on the left hemisphere were defined in AIBS space based on the AIBS atlas using 0.300 mm^3^ spheres, corresponding to 27 voxels (**Figure S1a**). Mean BOLD signal time series within a seed were extracted (*fslmeants*) and regressed into individual scans to obtain z-statistic maps (*fsl_glm*). Multi-session temporal concatenation ICA was carried out using MELODIC (Multivariate Exploratory Linear Optimized Decomposition into Independent Components, v3.14) using 20 components. Group-level component classification was adapted on a set of rules defined in (Zerbi et al., 2015). The following were considered plausible resting-state networks: (i) components with either bilateral organisation or (ii) unilateral components with a corresponding separate contralateral component, (iii) minimal crossing of relevant brain boundaries such as white matter tracts, (iv) spatial extent covering more than one slice. The following were considered as implausible resting-state networks: (i) components overlapping mainly with either white matter, ventricle, or vascular masks (**Supplementary figure 4bcd**), (ii) components mainly localised on brain edges. Dual-regression was carried out using the eponymous FSL function to obtain individual-level representations of 14 selected plausible group-level components (Filippini et al., 2009).

### Statistical analysis and data representation

Voxelwise statistics were carried out in FSL using either non-parametric permutation tests (*randomise*) for across datasets one-sample t-tests using 5000 permutations and voxelwise correction, or uncorrected parametric one-sample t-tests for within-dataset comparisons (*fsl_glm*). Voxelwise statistical maps are shown as colour-coded t-statistics overlays on the ABI template resampled at 25µm^3^ isotropic using MRIcron (Rorden et al., 2007). Statistical analysis carried out on parameters extracted from ROIs was performed in R (v3.4.4, “Someone to Lean on”, R Foundation for Statistical Computing, Vienna, Austria, https://R-project.org) using a linear model (*lm*). A simplified model was designed including the following fixed effects: breathing conditions (2 levels: ventilated or free-breathing), sedation conditions (4 levels: awake, isoflurane/halothane, medetomidine, medetomidine + isoflurane combination), SNR (continuous variable), mean FWD (continuous variable). Interactions effects between these factors were not modeled. Fixed effects significance was tested using likelihood ratio test. Scan parameter occurrence rates were assessed with Chi-square test (*chisq.test*). Residual analysis was performed with QQ-plots to inspect normal distribution, Tukey–Anscombe plots for the homogeneity of the variance and skewness, and scale location plots for homoscedasticity (i.e., the homogeneity of residual variance). The assumption of normality of the residuals was considered plausible in all statistical tests. Plots were generated using ggplot2 (v2.1.0) package for R. Significance level was set at p<=0.05 one-tailed with family-wise error correction at a voxelwise level, unless specified otherwise. Descriptive statistics are given as mean ± 1 standard deviation.

## Acknowledgements

This work was supported by the Singapore Bioimaging Consortium (SBIC), A*STAR, Singapore. AG acknowledges funding from the Simons Foundation (SFARI 314688 and 400101), the Brain and Behavior Foundation (2017 NARSAD independent Investigator Grant) and the European Research Council (ERC, G.A. 802371). This work was also supported by the JSPS KAKENHI Grant Number 16K07032 to NT, Brain/MINDS, the Strategic Research Program for Brain Sciences (SRPBS) from the Ministry of Education, Culture, Sports, Science, and Technology of Japan (MEXT) and Japan Agency for Medical Research and Development (AMED) to NT and HO. It was further supported as part of the Excellence Cluster ‘BrainLinks-BrainTools’ by the German Research Foundation, grant EXC1086. AH acknowledges funding from the German BMBF (NeuroImpa, 01EC1403C and NeuroRad 02NUK034D). MD acknowledges funding from France-Alzheimer Association, Plan Alzheimer Foundation and the French Public Investment Bank’s “ROMANE” program. This work was also supported by the Fund for Scientific Research Flanders (FWO) (grant agreements G057615N and 12S4815N -AvL), the Stichting Alzheimer Onderzoek (SAO-FRA, grant agreement 13026-AvL), the interdisciplinary PhD grant BOF DOCPRO 2014 -MV). The authors would like to thank Itamar Kahn, Eyal Bergmann and Daniel Gutierrez-Barragan for critically reading the manuscript.

## Author Contributions

JG designed the study. Every author contributed to data acquisition. JG, CC and AG carried out the analysis. Every author participated in the preparation of the manuscript.

## Competing interests statement

The authors have no conflicts of interest to declare.

