## Supplementary figures and tables for "Common functional networks in the mouse brain revealed by multi-centre resting-state fMRI analysis"

Contents: 17 Pages, 1 List of abreviations, 12 Figures, 3 Tables

**List of abbreviations**

| | |
|---|---|
| ACA | Anterior cingulate area |
| ACB | Nucleus accumbens |
| AI | Insular area |
| AIBS | Allen Institute for Brain Sciences |
| CP | Caudoputamen |
| DMN | Default-mode network |
| EPI | Echo planar imaging |
| FC | Functional connectivity |
| FWD | Framewise displacement |
| GSR | Global signal regression |
| HIP | Hippocampal region |
| ICA | Independent component analysis |
| MC | Motion parameter correction |
| MO | Somatomotor area |
| PAL | Pallidum |
| ROI | Region-of-interest |
| RSN | Resting state networks |
| RSP | Retrosplenial area |
| SBA | Seed-based analysis |
| SN | Salience networks |
| SNR | Signal-to-noise ratio |
| SSp | Primary somatosensory cortex |
| TH | Thalamus |
| tSNR | Temporal signal-to-noise ratio |
| VASC | Vascular |
| VEN | Ventricular |
| WM | White matter |

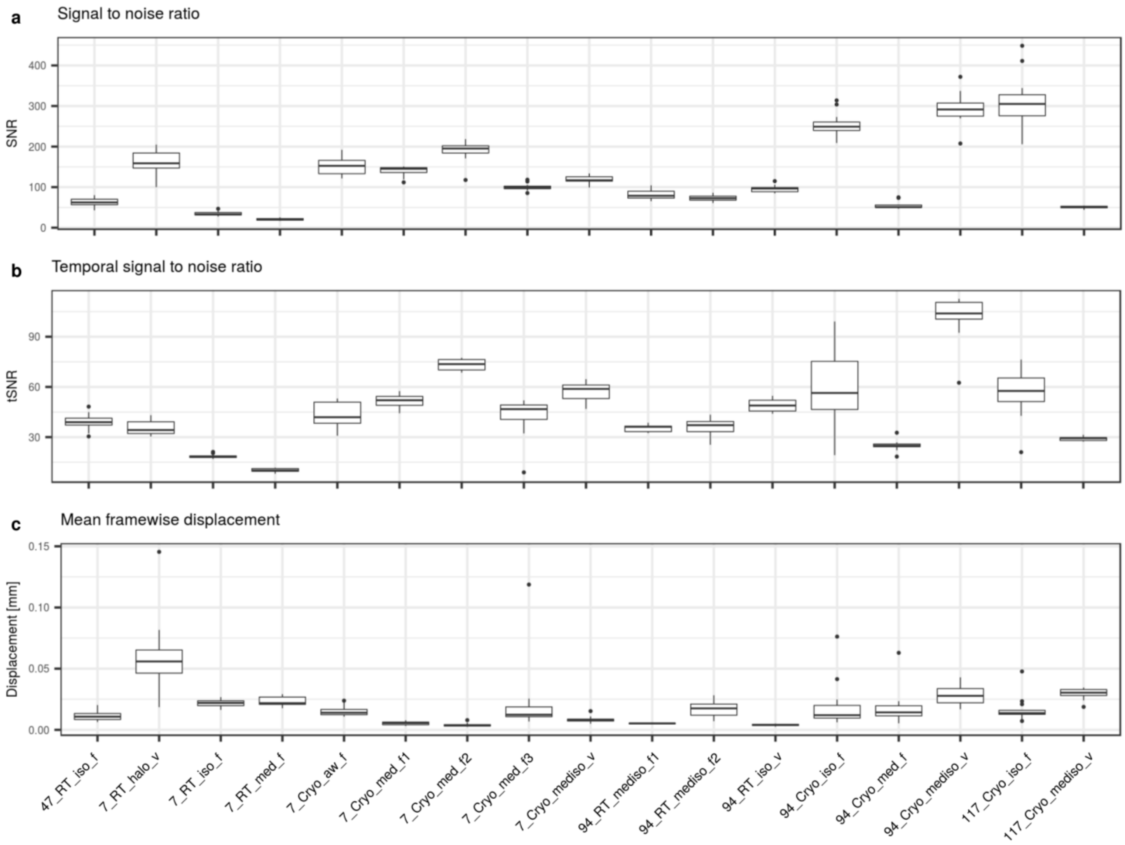


**Supplementary figure 1 |** Dataset description. Signal-to-Noise Ratio (SNR), Temporal SNR (tSNR), and mean framewise displacement as a function of dataset. There is a positive association between temporal SNR and SNR ($r = 0.75$, $t = 18.30$, $df = 253$ $p = 2.2e-16$).

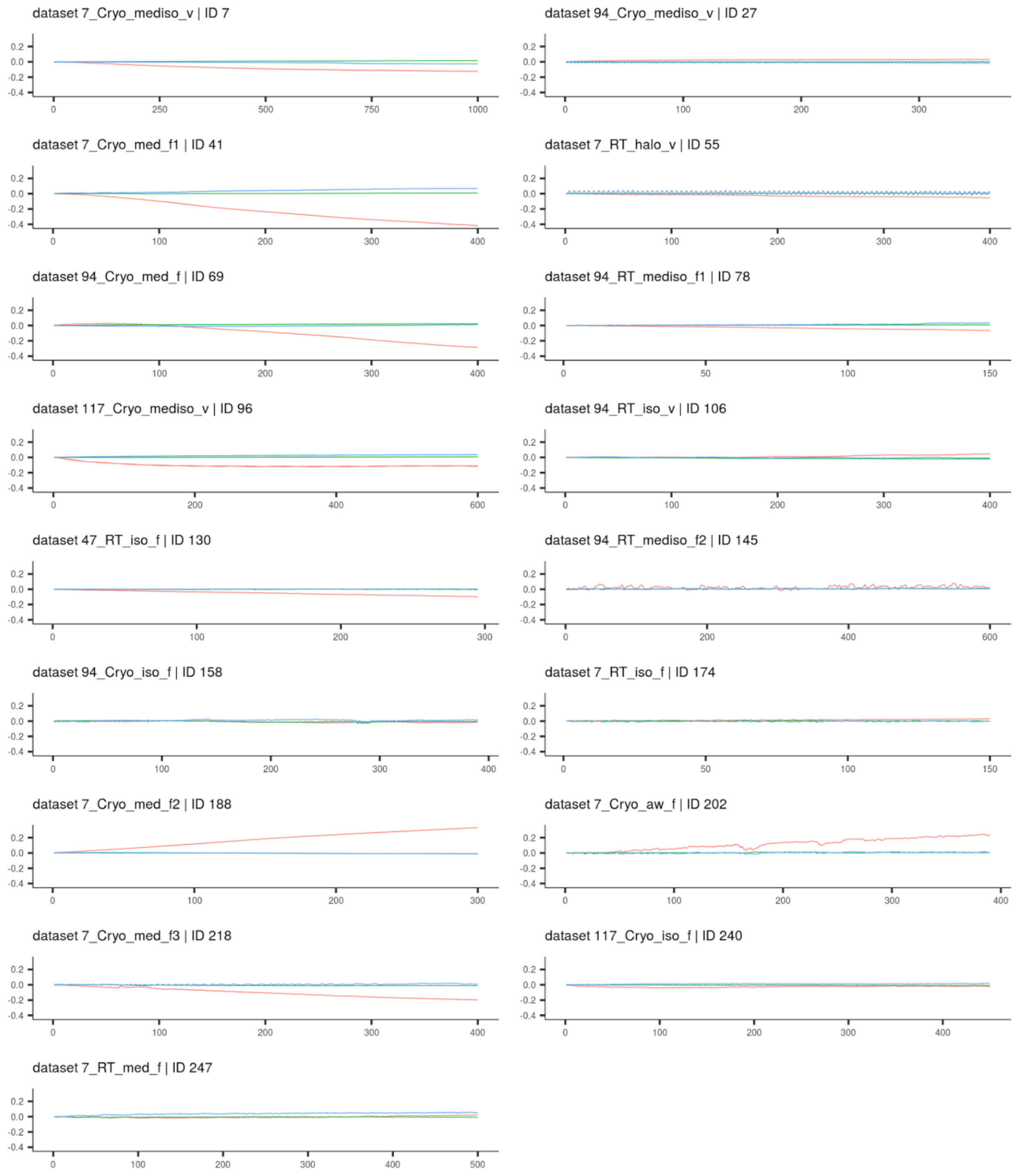


**Supplementary figure 2 |** Representative motion traces for one scan in each dataset. Nine datasets presented a visible continuous drift along the superior-inferior axis (dS, red) corresponding to the phase-encoding direction along the superior-inferior axis. Displacement is indicated in mm relative to the first volume.

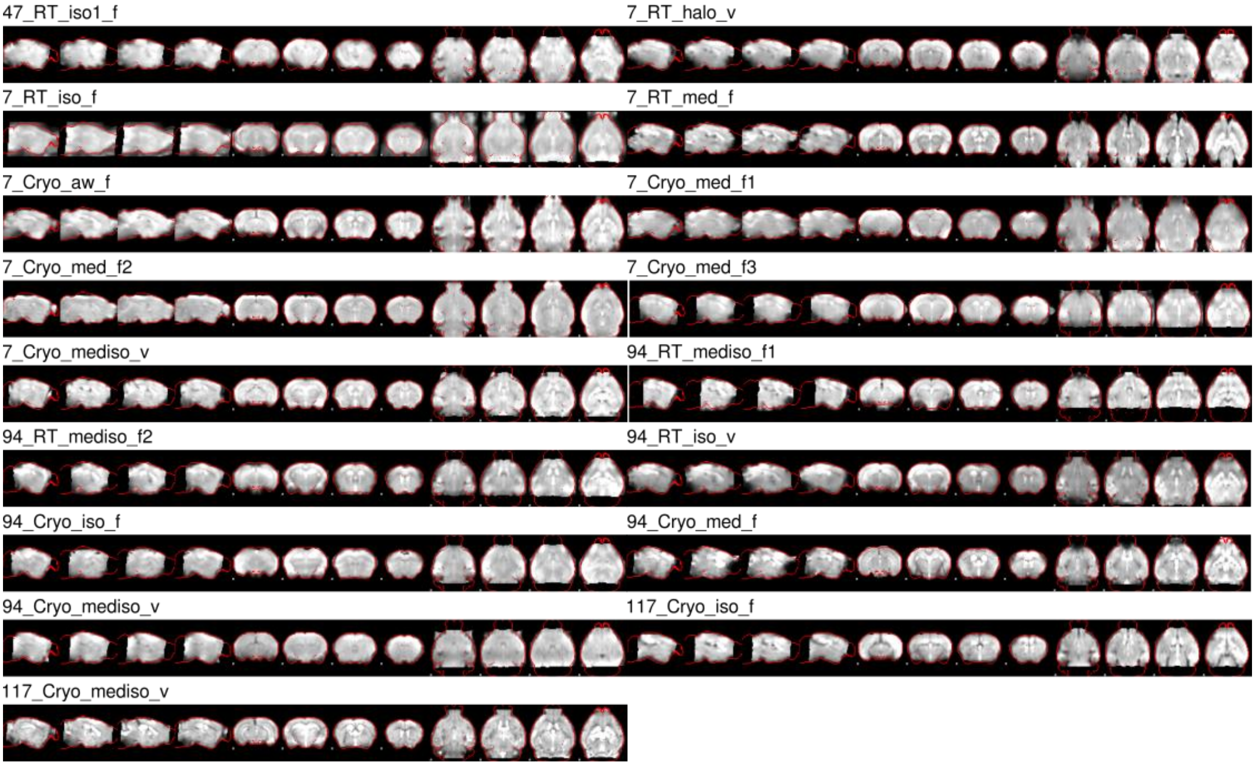


**Supplementary figure 3 |** Representative normalisation to AIBS template for each dataset showing 4 axial, 4 coronal, and 4 sagittal slices. Echo planar images were normalised to AIBS space (red outline) using a T2 weighted MRI template. Registrations and brain masks were considered plausible for all scans and datasets.

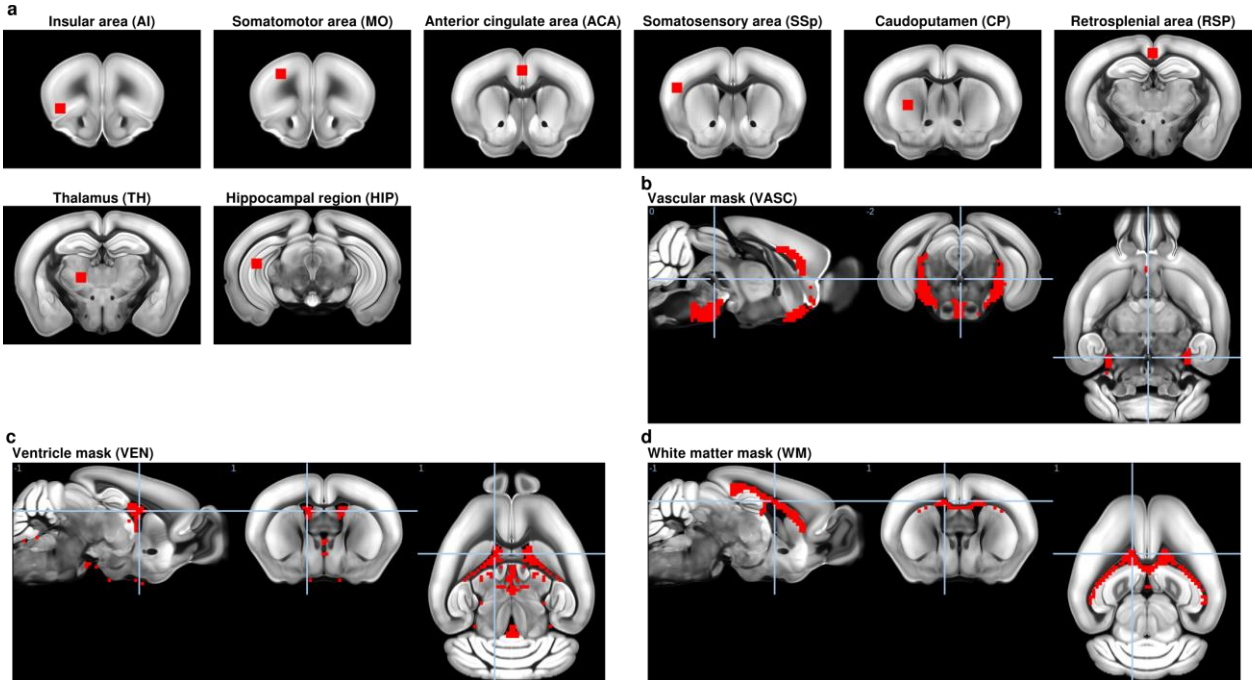


**Supplementary figure 4 |** Seed locations used in the present study (**a**). A data-driven vascular mask was derived from averaging selected independent components carried out on individual scans prior to applying denoising procedures (**b**). Ventricles (**c**) and white matter (**d**) masks were derived from the Allen Institute for Brain Science atlas and resampled to 200 $\mu m^3$ .

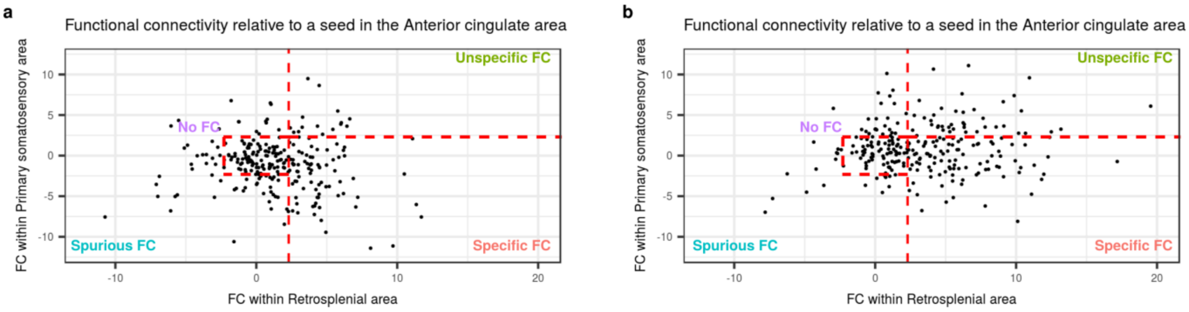


**Supplementary figure 5** | Individual-level FC parameters were extracted from the retrosplenial area (RSP, specific ROI) and primary somatosensory area (SSp, non-specific ROI) following (**a**) global signal regression and (**b**) vascular + ventricle signal regression. “Specific FC” is defined as high (z>=2.3, corresponding to p<0.01, one-tailed, uncorrected) FC in the RSP ROI and low (z<2.3) FC in the SSp. “Unspecific FC” is defined as high FC in both ROIs. “No FC” is defined as low absolute FC (abs(z)<2.3) in both ROIs. “Spurious FC” is defined as high absolute FC (abs(z)>=2.3) in non-specific ROI and low FC in specific ROI. 98/255 scans fell under the “specific FC” category for VV signal regression, the remaining presented “unspecific FC” (41/255), “spurious FC” (38/255), or “no FC” (78/255). GSR reduced FC within both the specific ROI and unspecific ROI, thus strengthening anti-correlation patterns in SSp relative to the anterior cingulate area.

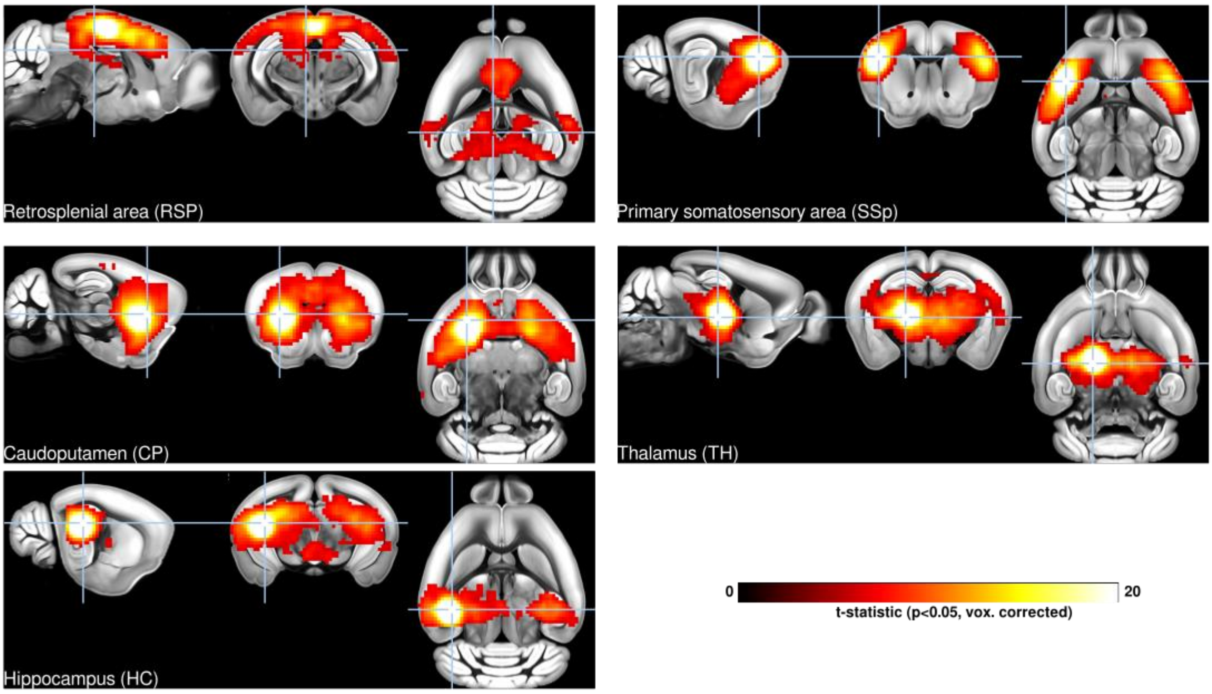


**Supplementary figure 6 |** SBA for 5 seeds positioned on the left hemisphere, including a seed in the retrosplenial area, primary somatosensory area, caudoputamen, hippocampus, and thalamus. Only SBA-derived from 98/255 scans that presented "specific FC" following vascular+ventricle signal regression were included. All FC maps reveal a mainly bilateral homotopic organisation. These 5 seeds complement the 3 seeds presented in **Figure 2**. The corresponding one-sample t-test including 255/255 scans in shown in **Supplementary figure 7**. Voxelwise corrected t-statistic for one-sample t-tests are shown as a colour-coded overlay on the AIBS reference template.

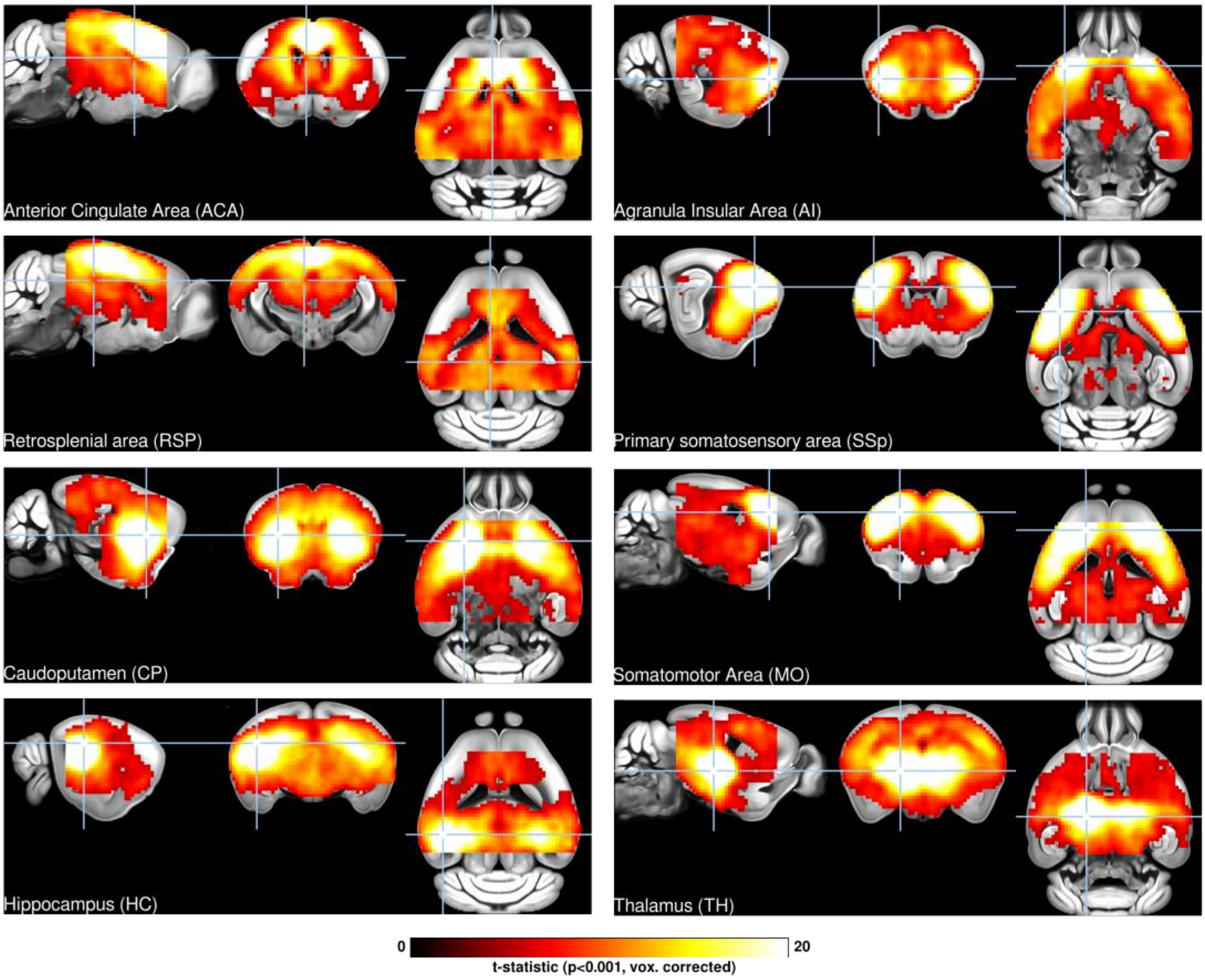


**Supplementary figure 7** | One-sample t-test of SBA for 8 seeds examined in 255/255 scans. Voxelwise corrected t-statistic for one-sample t-tests ($p<0.001$, corrected) are shown as a colour-coded overlay on the AIBS reference template.

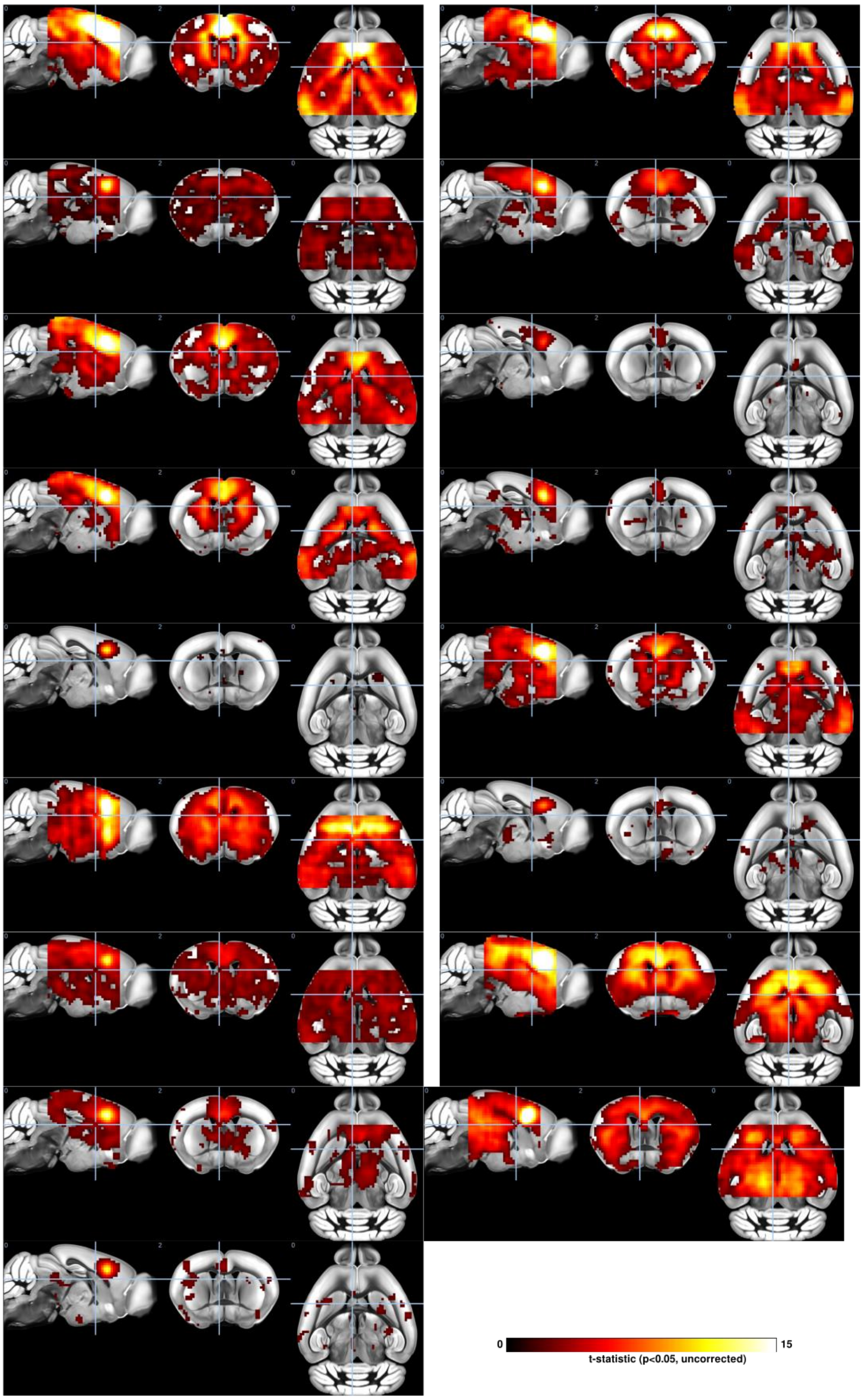


**Supplementary figure 8 |** SBA of the DMN derived from individual dataset, each consisting of 15 scans. Uncorrected t-statistic for one-sample t-tests (p<0.05, uncorrected) are shown as a colour-coded overlay on the AIBS reference template.

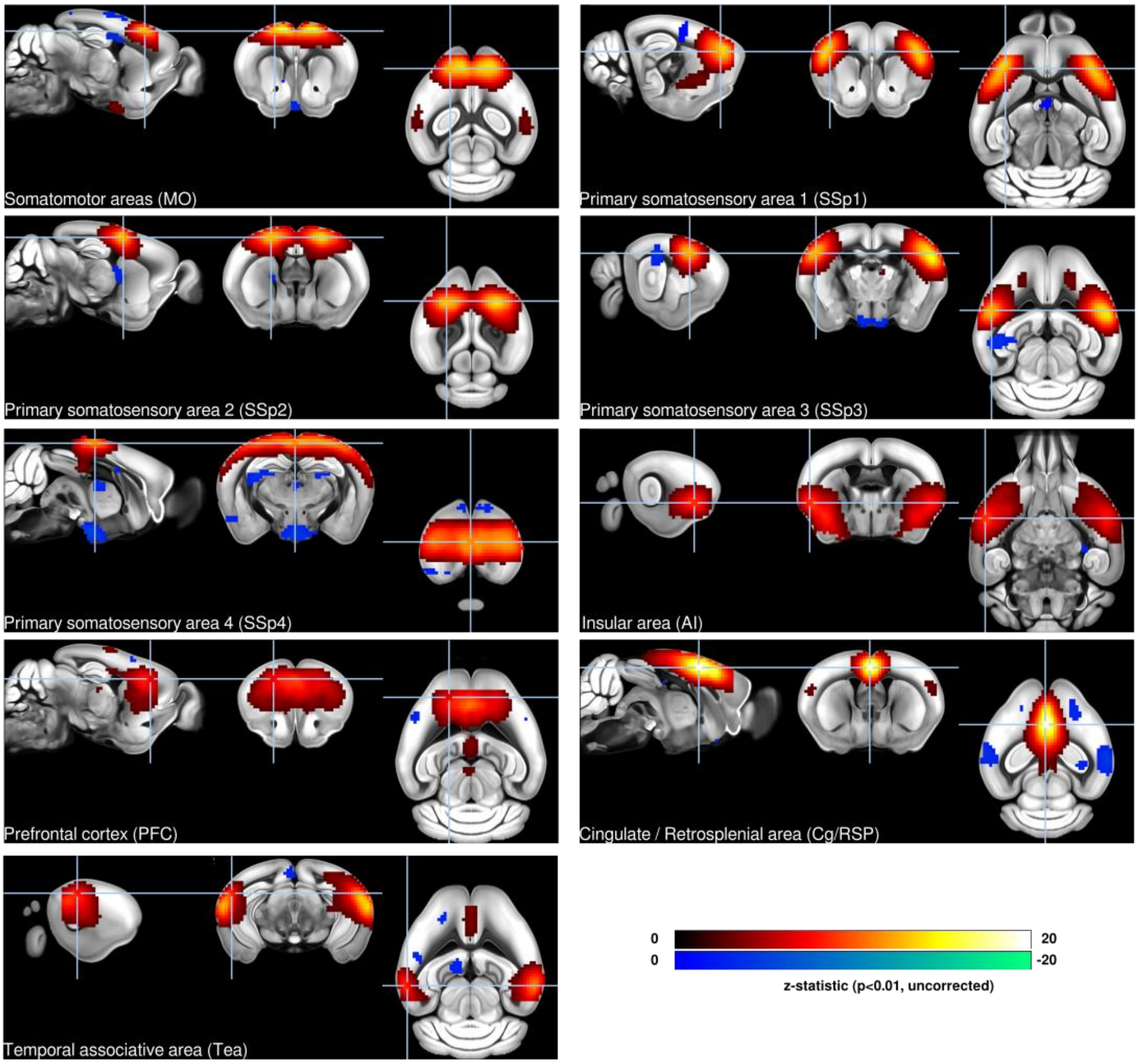


**Supplementary figure 9 |** Details of the 9 cortical components detected by group ICA. Only 99/255 scans that presented "specific FC" following vascular+ventricle signal regression were included in the group ICA estimation.

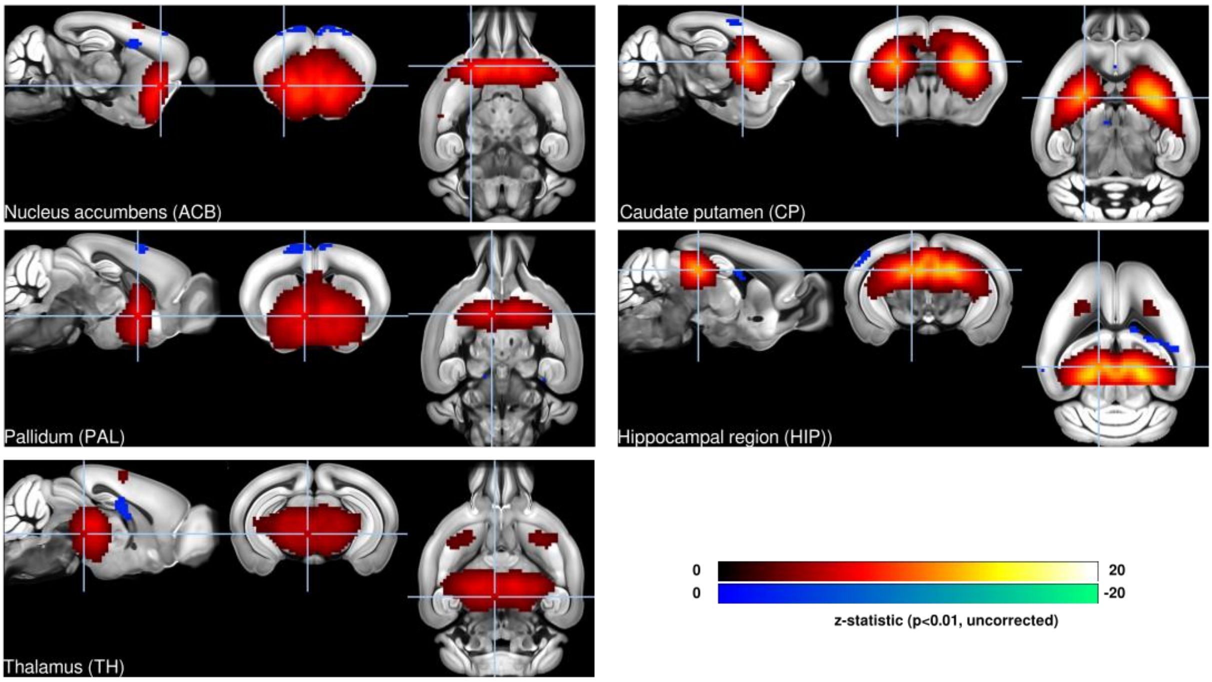


**Supplementary figure 10 |** Details of the 5 sub-cortical components detected by group ICA. Only 98/255 scans that presented “specific FC” following vascular+ventricle signal regression were included in the group ICA estimation.

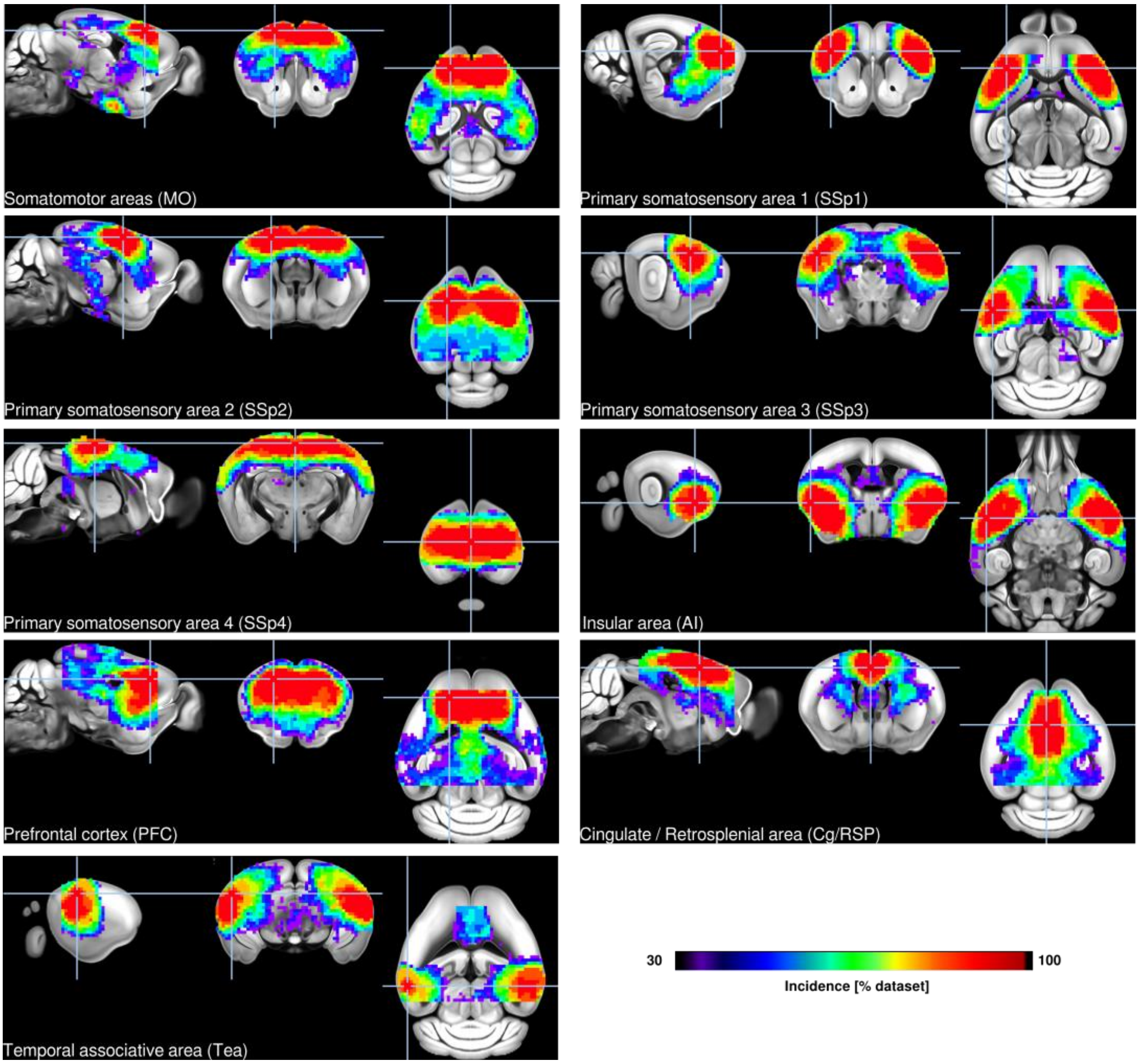


**Supplementary figure 11 |** Incidence maps representing the percentage of dataset surviving the significance threshold in a parametric one-sample t-test ($p < 0.05$, uncorrected) following a dual-regression analysis using 9 cortical independent components as spatial references (**Figure 4**). Incidence reached 100% within the main component clusters in all cases, denoting that significant FC could be established in all datasets.

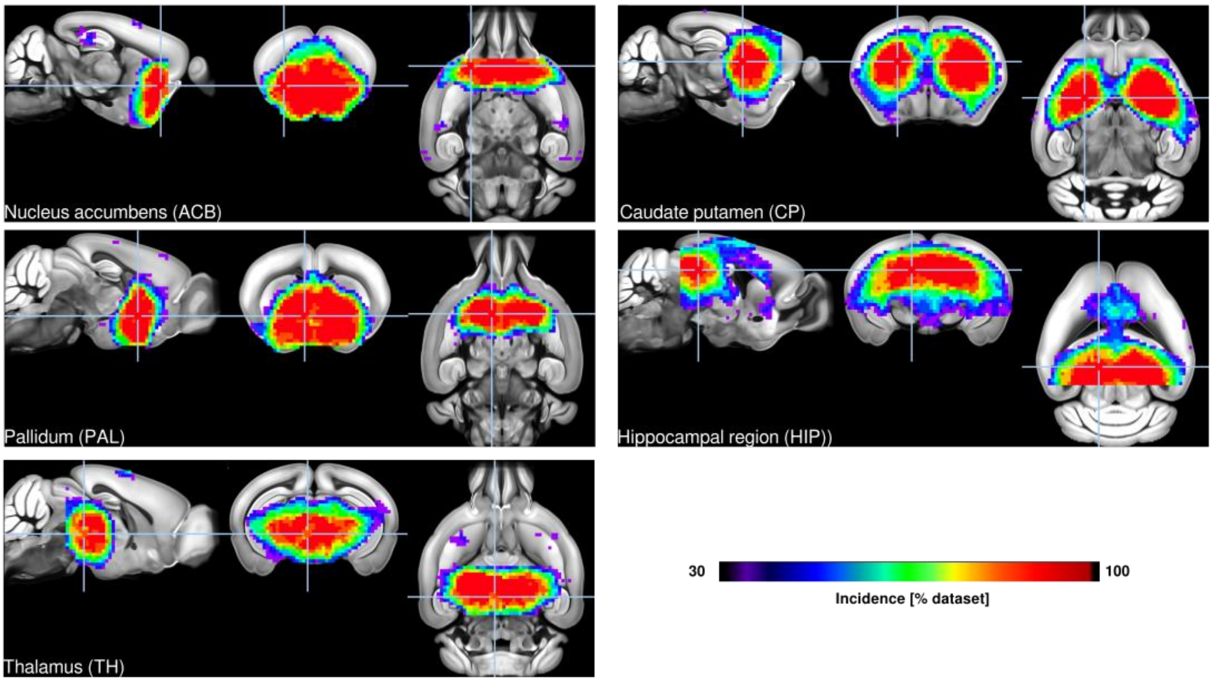


**Supplementary figure 12 |** Incidence maps representing the percentage of dataset surviving the significance threshold in a parametric one-sample t-test ($p < 0.05$, uncorrected) following a dual-regression analysis using 5 sub-cortical independent components as spatial references (**Figure 4**). Incidence reached 100% within the main component clusters in all cases, denoting that significant FC could be established in all datasets.

### Tables

| Dataset ID | Equipment | EPI parameters | Animal handling | Mean FWD [mm] | SNR | tSNR | Excluded frames (%) |
|---|---|---|---|---|---|---|---|
| 47_RT_iso1_f | 4.7T<br>2x2 array (R) | TR/TE 2000/25.03<br>FOV 15x15<br>Mat 64x64<br>Slices/Thickness 22/0.5<br>Volumes 295 | free-breathing<br>Isoflurane 1%<br>Breath per min 100 | 0.011±0.004 | 62.4±9.9 | 39.2±4.6 | 0 |
| 7_RT_halo_v | 7T<br>2 x 2 array (R) | TR/TE 1200/15<br>FOV 20x20<br>Mat 100x100<br>Slices/Thickness 18/0.5<br>Volumes 400 | Ventilated<br>Halothane 0.75%<br>Breath per min 90 | 0.060±0.028 | 161±28.4 | 35.8±4 | 10.15±20.63 |
| 7_RT_iso_f | 7T<br>23mm Volume coil (T/R) | TR/TE 2000/15<br>FOV 19.2x9<br>Mat 64x30<br>Slices/Thickness 15/0.75<br>Volumes 150 | Free-breathing<br>Isoflurane 1%<br>Breath per min 100 | 0.022±0.003 | 34.4±5.2 | 18.5±1.2 | 0 |
| 7_RT_med_f | 7T<br>Surface coil (T/R) | TR/TE 2000/15<br>FOV 21.2x18.46mm<br>Mat 147x59<br>Slices/Thickness 31/0.5<br>Volumes 500 | free-breathing<br>Medetomidine 0.3 mg/kg \| 0.6 mg/kg/h (s.c.)<br>Breath per min NA | 0.023±0.004 | 20.6±2.5 | 10.2±1.1 | 0 |
| 7_Cryo_aw_f | 7T<br>Cryoprobe surface (T/R) | TR/TE 1500/15<br>FOV 12.8x12.8mm<br>Mat 64x64<br>Slices/Thickness 18/0.75<br>Volumes 390 | free-breathing<br>Awake<br>Breath per min NA | 0.015±0.004 | 150.1±21.4 | 43.3±7.5 | 0.38±0.68 |
| 7_Cryo_med_f1 | 7T<br>Cryoprobe surface (T/R) | TR/TE 1700/10<br>FOV 19.5x12mm<br>Mat 128x80<br>Slices/Thickness 22/0.7<br>Volumes 400 | free-breathing<br>Medetomidine 0.3 mg/kg \| 0.6 mg/kg/h (s.c.)<br>Breath per min 115 | 0.005±0.001 | 140.2±11.8 | 51.8±3.7 | 0.03±0.09 |
| 7_Cryo_med_f2 | 7T<br>Cryoprobe surface (T/R) | TR/TE 2000/10<br>FOV 19.2x12mm<br>Mat 128x80<br>Slices/Thickness 30/0.5<br>Volumes 300 | free-breathing<br>Medetomidine 0.3 mg/kg \| 0.6 mg/kg/h (s.c.)<br>Breath per min 70-100 | 0.004±0.001 | 190.1±24 | 73.2±3.4 | 0 |
| 7_Cryo_med_f3 | 7T<br>Cryoprobe surface (T/R) | TR/TE 1700/10<br>FOV 19.2x12mm<br>Mat 128x80<br>Slices/Thickness 12/0.7<br>Volumes 400 | free-breathing<br>Medetomidine 0.3 mg/kg \| 0.6 mg/kg/h (s.c.)<br>Breath per min 115 | 0.021±0.028 | 100.3±8.1 | 42.8±11.1 | 2.57±9.87 |
| 7_Cryo_mediso_v | 7T<br>Cryoprobe surface (T/R) | TR/TE 1000/15<br>FOV 20x10<br>Mat 90x50<br>Slices/Thickness 20/0.5<br>Volumes 1000 | Ventilated<br>Isoflurane 0.5% + Medetomidine 0.05mg/kg \| 0.1mg/kg/h (i.v.)<br>Breath per min 80 | 0.008±0.002 | 119.4±9.5 | 57.5±5.2 | 0.13±0.44 |
| 94_RT_mediso_f1 | 9.4T<br>Quadrature surface (R) | TR/TE 2000/15<br>FOV 20x20<br>Mat 128x64<br>Slices/Thickness | Free-breathing<br>Isoflurane 0.4% + Medetomidine 0.3mg/kg (s.c) | 0.005±0.000 | 81.5±11.1 | 35.4±2 | 0 |

| | | | | | | | |
|---|---|---|---|---|---|---|---|
| | | 16/0.5<br>Volumes 150 | Breath per min NA | | | | |
| 94_RT_mediso_f2 | 9.4T<br>2x2 array (R) | TR/TE 1000/15<br>FOV 20x20<br>Mat 64x64<br>Slices/Thickness 16/0.5<br>Volumes 600 | Free-breathing<br>Isoflurane 0.3% +<br>Medetomidine 0.05mg/kg \| 0.1mg/kg/h (s.c.)<br>Breath per min 120 | 0.017±0.006 | 73.1±6.8 | 35.6±5.6 | 0.14±0.27 |
| 94_RT_iso_v | 9.4T<br>2x2 array (R) | TR/TE 1500/17<br>FOV 17x9<br>Mat 70x35<br>Slices/Thickness 18/0.6<br>Volumes 400 | Ventilated<br>Isoflurane 1%<br>Breath per min 80 | 0.004±0.001 | 95.6±8 | 48.8±3.5 | 0 |
| 94_Cryo_iso_f | 9.4T<br>Cryoprobe surface (T/R) | TR/TE 1000/10<br>FOV 23.7×14<br>Mat 90×60<br>Slices/Thickness x/0.45<br>Volumes 400 | free-breathing,<br>Isoflurane 1.2%<br>Breath per min 100 | 0.020±0.018 | 251.4±29.6 | 59.8±22.3 | 2.43±7.39 |
| 94_Cryo_med_f | 9.4T<br>Cryoprobe surface (T/R) | TR/TE 1300/18<br>FOV 17.28×11.52<br>Mat 96×64<br>Slices/Thickness 21/0.4<br>Volumes 400 | free-breathing<br>Medetomidine 0.4 mg/kg \| 0.8 mg/kg/h (i.p)<br>Breath per min NA | 0.018±0.014 | 54.5±8.4 | 24.9±3 | 0.72±2.71 |
| 94_Cryo_mediso_v | 9.4T<br>2x2 Cryoprobe array (R) | TR/TE 1000/9.2<br>FOV 20x17.5<br>Mat 90x70<br>Slices/Thickness 12/0.5<br>Volumes 360 | Ventilated<br>Isoflurane 0.5% +<br>Medetomidine 0.05mg/kg \| 0.1mg/kg/h (i.v.)<br>Breath per min 80 | 0.027±0.008 | 293.3±37 | 102.5±12.7 | 0 |
| 117_Cryo_iso_f | 11.7T<br>Cryoprobe surface (T/R) | TR/TE 1000/10<br>FOV 22.5x15.4<br>Mat 90x70<br>Slices/Thickness 12/0.7<br>Volumes 450 | Free-breathing,<br>Isoflurane 1.25%<br>Breath per min 100 | 0.016±0.010 | 309.7±60.1 | 57.1±13.6 | 0.18±0.69 |
| 117_Cryo_mediso_v | 11.7T<br>2x2 Cryoprobe array (R) | TR/TE 1000/15<br>FOV 17x9<br>Mat 90x60<br>Slices/Thickness 28/0.4<br>Volumes 600 | Ventilated<br>Isoflurane 0.5% +<br>Medetomidine 0.05mg/kg \| 0.1mg/kg/h (s.c.)<br>Breath per min 80 | 0.030±0.004 | 50.6±2.7 | 29.1±1.2 | 0.06±0.14 |

**Supplementary table 1 |** Dataset description

| ROI pair | **Restraint** $F_{(247,251)}$, **p** | **Breathing** $F_{(247,248)}$, **p** | **Motion** $F_{(247,248)}$, **p** | **SNR** $F_{(247,248)}$, **p** | **Percentile** $50^{th}$, $75^{th}$, $95^{th}$ |
|---|---|---|---|---|---|
| ACA - RSP | **18.02, 5.3e-13** | **3.48, 6.34e-2** | 0.085, 7.7e-1 | **12.59, 4.7e-4** | 2.70, 5.66, 10.46 |
| AI left - AI right | **27.74, 4.9e-19** | 0.042, 8.4e-1 | 0.098, 7.5e-1 | **87.33, 5.7e-18** | 4.18, 8.90, 15.65 |

| | | | | | |
|---|---|---|---|---|---|
| MO left - MO right | **15.99, 1.2e-11** | 5.97, 1.5e-02 | 0.40, 5.3e-1 | **66.68, 1.7e-14** | 5.84, 12.06, 20.84 |
| SSp left - SSp right | **10.69, 5.2e-8** | **16.85, 5.5e-5** | 0.97, 3.3e-1 | **64.19, 4.5e-15** | 5.23, 12.03, 19.45 |
| CP left - CP right | **13.97, 2.7e-10** | 0.71, 4.0e-1 | **4.32, 3.9e-2** | **100.68, 4.3e-20** | 4.53, 9.71, 16.39 |
| HIP left - HIP right | **17.72, 8-41e-13** | **12.05, 6.1e-4** | 0.54, 4.6e-1 | **89.28, 2.8e-18** | 2.35, 6.13, 15.70 |
| TH left - TH right | **24.79, 2.9e-17** | **4.62, 3.3e-2** | 0.61, 4.3e-1 | **135.47, 2.98e-25** | 2.88, 7.40, 19.37 |

**Supplementary table 2 |** Fixed effect analysis and functional connectivity (z-statistics) distribution as function of ROI pairs

**Supplementary table 3 |** Independent component functional connectivity strength (z-statistics) as a function of the Allen Brain Institute atlas.
